# Enhanced microvasculature formation and patterning in iPSC–derived kidney organoids cultured in physiological hypoxia

**DOI:** 10.1101/2021.12.22.473849

**Authors:** A. Schumacher, N. Roumans, T. Rademakers, V. Joris, M. Eischen-Loges, M. van Griensven, V.L.S. LaPointe

## Abstract

Functional kidney organoids have the potential to be used in implantable kidney grafts for patients with end-stage kidney disease, because they have been shown to self-organize from induced pluripotent stem cells into most important renal structures. To date, however, long-term kidney organoid culture has not succeeded, as nephrons lose their phenotype after approximately 25 days. Furthermore, the renal structures remain immature with diminishing endothelial networks with low connectivity and limited organoid invasion. We hypothesized that introducing long-term culture at physiological hypoxia, rather than the normally applied non-physiological, hyperoxic 21% O_2_, could initiate angiogenesis, lead to enhanced growth factor expression and improve the endothelial patterning. We therefore cultured the kidney organoids at 7% O_2_ instead of 21% O_2_ for up to 25 days and evaluated nephrogenesis, VEGF-A expression and vascularization. Whole mount imaging revealed a homogenous morphology of the endothelial network with enhanced sprouting and interconnectivity when the kidney organoids were cultured in hypoxia. Three-dimensional quantification confirmed that the hypoxic culture led to an increased average vessel length, likely due to the observed upregulation of proangiogenic *VEGF-A189* mRNA and downregulation of the antiangiogenic protein VEGF-A165b measured in hypoxia. This research indicates the importance of optimization of oxygen availability in organoid systems and the potential of hypoxic culture conditions in improving the vascularization of organoids.

**Significance statement:** Culturing kidney organoids in a hypoxic environment improved patterning of the endothelial network and improved vascularization. These findings may help improve the quality of kidney organoids, and could eventually improve the kidney graft for transplantation in patients with end-stage kidney disease. Furthermore, the organoids will be more suitable for drug testing and in developmental biology. The findings might also be translatable to other organoid models containing endothelial cells.

**Graphical abstract:** 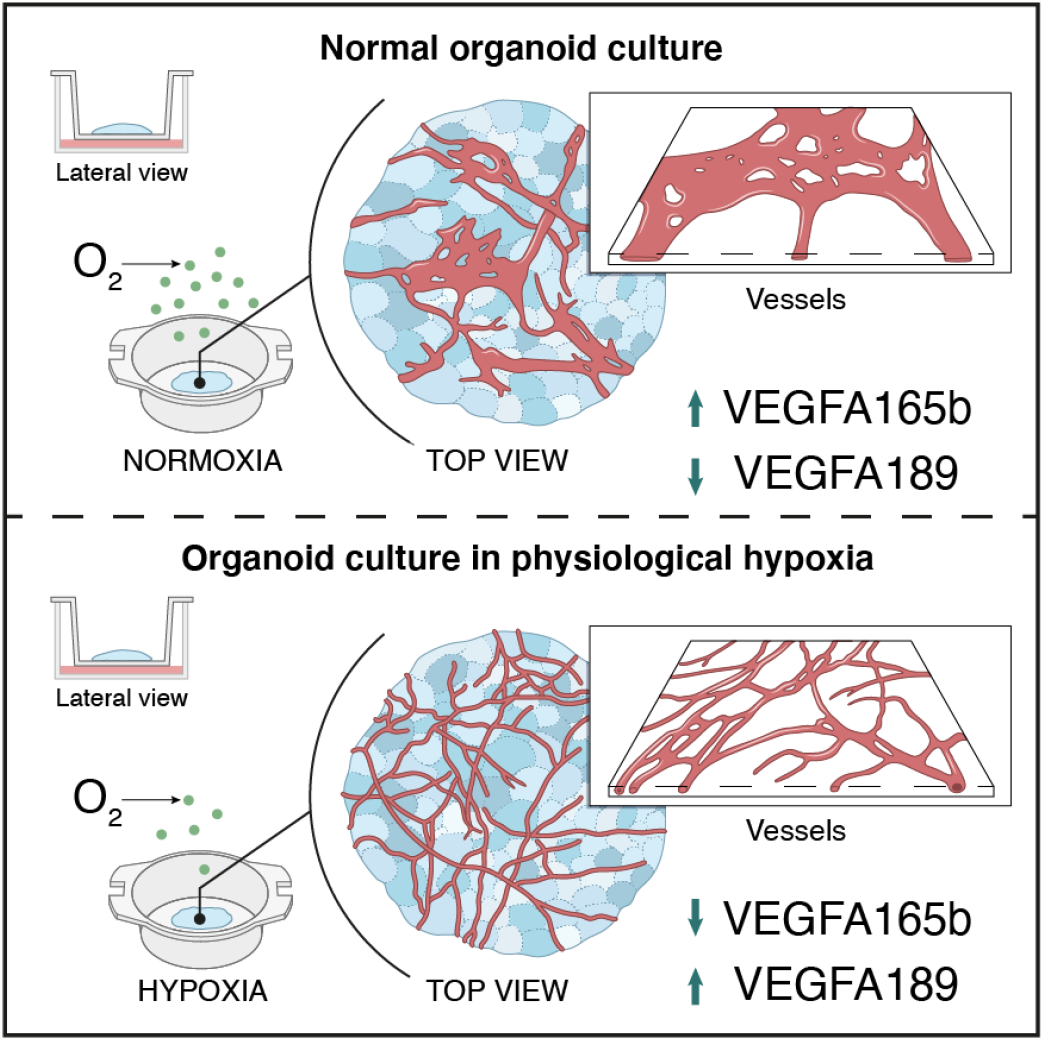

## 1 Introduction

Organoid models have become an irreplaceable alternative to two-dimensional cell culture because of their greater cellular and architectural complexity, which is alike native tissue. Induced pluripotent stem cells can be differentiated into kidney organoids that develop nephrons resembling capillary loop–stage nephrons.^1^ The large variety of renal cell types they possess, similar to the developing kidney^2^, makes them promising models for regenerative medicine, drug testing and developmental biology^3^. However, these organoids have several drawbacks—such as the limited culture duration, loss of nephrogenic potential, immaturity and lack of vasculature^4,5^—perhaps due the lack of an *in vivo*–like culture environment, which is increasingly being investigated.^6^ Moreover, kidney organoids are one of the largest organoid models, growing up to 1.5 mm in thickness.^1^ This draws attention to one aspect of a physiological environment, which is the oxygenation of the tissue.

Cell culture in hypoxia, defined as a state where cells no longer have sufficient oxygen available to use oxidative phosphorylation to generate ATP^7^, has been performed for more than a decade, for example to maintain stem cells^8^. Hypoxia is also known to act as a morphogen in cell communication of various cell lineages and for certain cell types to partially determine cellular differentiation.^9^ If not optimized to the model system, hypoxia is known to have detrimental effects in cell and tissue culture. Nevertheless, to date, there is little knowledge on the effects of hypoxia in organoid cultures. Briefly, hypoxic culture (5% O_2_) of intestinal organoids was considered detrimental due to a reduced number of crypts.^10^ In contrast, microwell-cultured kidney organoids cultured for 24 hours in hypoxia (1% and 3% O_2_) showed enhanced functionality measured by secretion of erythropoietin (EPO).^11^ These results argue that oxygen levels critically affect maturation of organoids, in a model-sensitive manner.

Kidney organoids are cultured, like explanted kidneys^12^, at an air–liquid interface and therefore are directly exposed to incubator air (21% O_2_), commonly defined as normoxia. However, it has been previously well described that 21% O_2_ is non-physiological.^7^ For this reason, and the fact that the physiological environment of the developing kidney is hypoxic, we do consider 21% O_2_ to be hyperoxic and non-physiological. Hyperoxia is known to negatively impact kidney development *in vivo*, such as significantly reducing the size of the nephrogenic zone and glomeruli.^13^ *In vitro*, the detrimental effects of hyperoxic cell and tissue culture are also well established^14,15^, such as the formation of reactive oxygen species in endothelial cells^16^. By contrast, there is significant evidence that hypoxia enhances the proliferation of endothelial cells as well as stem cell differentiation towards an endothelial lineage.^17-23^ Murine metanephric explants showed enhanced endothelial cell proliferation when cultured in hypoxia (3% O_2_).^24,25^ Therefore, modulating the oxygen concentrations could be a step towards *in vivo*–like kidney organoid culture with improved vascularization.

In developing kidneys in the hypoxic mammalian uterus^26^, nephrogenesis starts in the avascular nephrogenic zone.^27^ Later in the capillary loop stage of nephrogenesis, mainly angiogenesis and to a lesser extent vascularization start to take place.^4^ Only when blood vessels enter the kidney and new vessels are formed, oxygen levels increase to finally reach 4– 9.5 kPa (30–71 mmHg) in the adult human cortex and 2 kPa (15 mmHg) in the adult human medulla.^28^ Due to the invasive nature of the measurements, the oxygen tensions in developing human cortex and medulla have not been determined. Recapitulating this *in vivo* hypoxic environment in the kidney organoid culture could enable the transcription of genes essential in kidney organogenesis. Below 5% O_2_, binding of prolyl hydroxylases to cytoplasmic hypoxia-inducible factor alpha (HIFα) is inhibited, leading to reduced or inhibited proteosomal degradation. Consequently, HIFα rapidly accumulates, translocates into the nucleus and its dimerization with HIFβ is initiated.^27^ Dimer binding to hypoxia responsive element (HRE) promoters leads to the transcription of a variety of genes. While this process occurs in all tissue types, it is known that various HIF-regulated genes are implicated in angiogenesis and vascularization in kidney organogenesis. In kidney development, nuclear HIF translocation occurs in metanephric mesenchyme and subsequently in podocytes, leading to VEGF-A transcription, which attracts endothelial cells to vascularize the nephrons.^29,30^

Developmentally, kidney organoids, normally cultured at non-physiological hyperoxic 21% O_2_ (≈160 mmHg), are comparable to kidneys in the first to early second trimester.^31^ We hypothesized whether a culture at 7% O_2_ (≈53 mmHg), comparable to the adult human cortex, would lead to intra-organoid oxygen concentrations more closely resembling developing non-vascularized kidneys *in vivo*, and initiate angiogenesis in the organoids. After 20 days of culture, we analyzed nephrogenesis, the expression of angiogenic markers (particularly VEGF) and endothelialisation to determine whether hypoxic cultures of the organoids improved these characteristics compared to cultures at 21% O_2_. We found that long-term hypoxic culture enhances endothelial patterning and sprouting and consequently could enhance kidney organoid cultures towards a more *in vivo*–like model.

## 2 Materials and Methods

### 2.1 Induced pluripotent stem cell differentiation and kidney organoid culture

Induced pluripotent stem cells (iPSCs) were differentiated and the organoids were cultured according to the previously published protocol by van den Berg, et al. ^32^ (Figure 1A). Briefly, the iPSC line LUMC0072iCTRL01 (male fibroblasts reprogrammed using RNA Simplicon reprogramming kit, Millipore) obtained from the hiPSC core facility at the Leiden University Medical Center, were passaged biweekly and maintained in 3 mL of Essential 8 Medium (Thermo Fisher Scientific) supplemented with 1% penicillin–streptomycin in 6-well plates coated with truncated recombinant human vitronectin (Thermo Fisher Scientific). The iPSC line was previously assessed for pluripotency and normal karyotype.^33^ For differentiation, 80,000 cells per well were seeded into a six well plate and differentiated according to the published protocol in organoid culture medium composed of STEMdiff APEL2 medium (STEMCELL Technologies), supplemented with 1% antibiotic-antimycotic and 1% protein-free hybridoma medium II (PFHM 2) (Thermo Fisher Scientific) and the respective growth factors and small molecules (8 µM GSK-3 inhibitor CHIR99021 (R&D Systems), 200 ng/mL FGF9 (fibroblast growth factor 9; R&D Systems), and 1 μg/mL heparin (Sigma-Aldrich)). After 7 days of differentiation, the cells were aggregated by centrifugation and spotted on transwell tissue culture plates with 0.4 μm pore polyester membrane inserts (Corning) and cultured at an air–liquid interface. For an additional 5 days, denoted as day 7+5, the organoids were cultured in organoid medium containing the growth factors and small molecules. From day 7+5 onwards, the organoids were cultured in organoid culture medium either in a hypoxia incubator (37°C, 5% CO_2_, 7% O_2_) or in normoxia (37°C, 5% CO_2_, 21% O_2_). The medium was refreshed every two days. At day 7+18 and day 7+25, the organoids were assessed.

**Figure 1.**
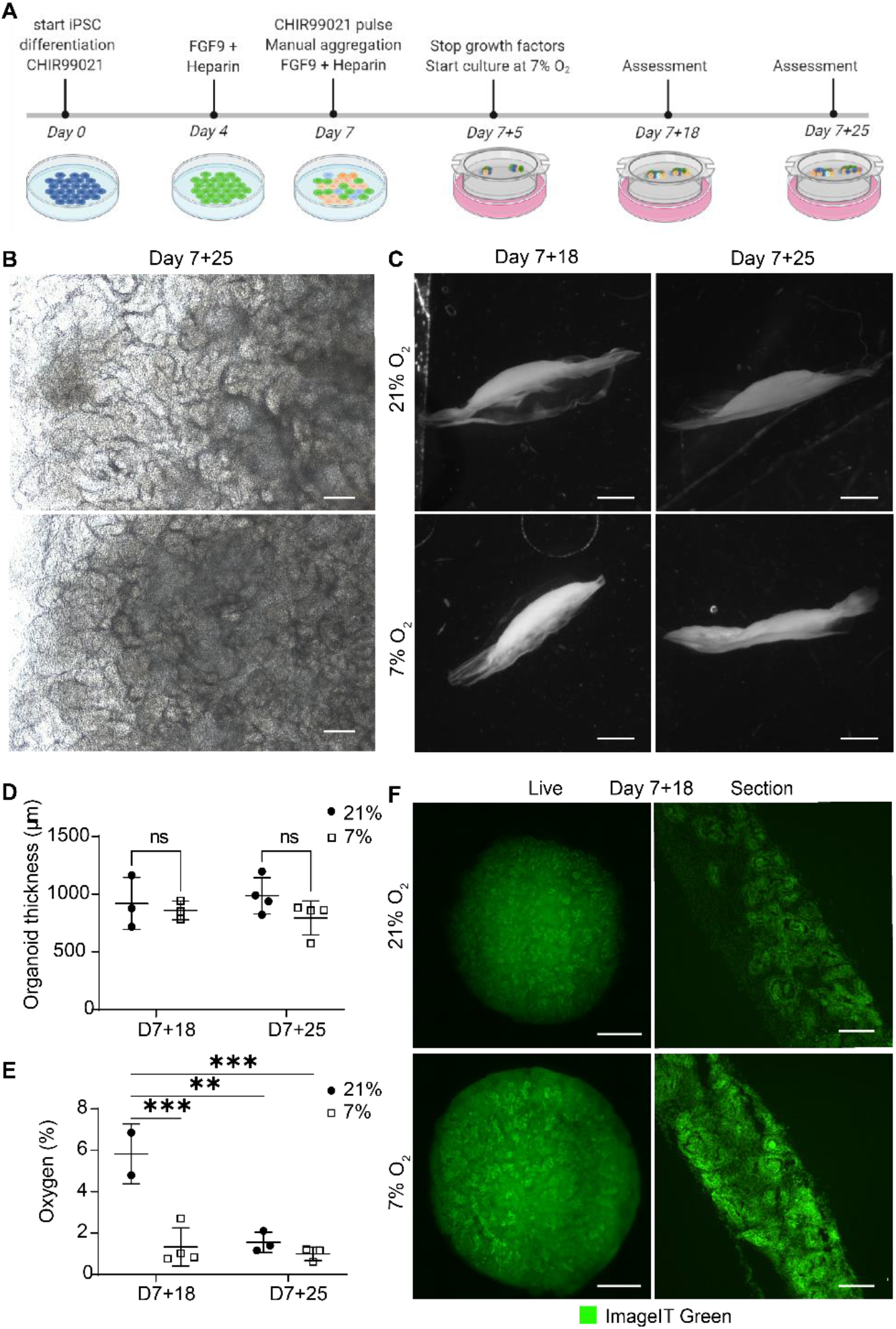
Kidney organoids cultured in uterine-like hypoxic/low oxygen environment structurally form like their normoxic counterparts without the appearance of a hypoxic core. A. Schematic representation of the iPSC differentiation and organoid culture. Human iPSCs are differentiated for 7 days, after which aggregates are spotted on the air–liquid interface and cultured as organoids for up to 25 additional days (termed day 7+25). B. Brightfield images display comparable morphologies of the organoids cultured in 7% and 21% O_2_ at the culture endpoint day 7+25. Scale bar: 200 µm. C. Darkfield images show that the macro-anatomy of the organoids is comparable in both oxygen concentrations. Scale bar: 2 mm. D. Quantification of the organoid thickness from the darkfield images shows no difference between 21% O_2_ and 7% O_2_. (n=2, N=3). E. Oxygen measurement in the bottom part of the organoids towards the transwell filter shows that organoids in 7% oxygen have a comparable oxygen concentration to their normoxic counterparts at day 7+25, which is significantly lower compared to the normoxic condition at day 7+18. (n=2, N=2; ** = *p* ≤ 0.01, *** = *p* ≤ 0.001) F. Organoids (left column) and cryosections of their central core (right column) stained on day 7+18 with the hypoxia dye ImageIT Green that produces a fluorescent signal below 5% O_2_. The organoids display no hypoxic core but the dye stains all nephrons. Scale bar “ live” : 1 mm, scale bar “ section” : 100 µm.

### 2.2 Darkfield imaging

The organoids were fixed at day 7+18 and day 7+25 in 2% (v/v) paraformaldehyde (20 min, 4°C) and were subsequently embedded in 10% gelatin in PBS (phosphate-buffered saline) in a cryomold. Once hardened, the gel was pealed out of the mold and placed on a darkfield imaging stage with circular illumination (Nikon). Images were acquired using a 1× air objective on a Nikon SMX25 automated stereomicroscope with a customized Nikon darkfield illumination holder.

### 2.3 Cryopreservation and cryosectioning

After fixing the organoids at day 7+18 or day 7+25 in 2% (v/v) paraformaldehyde (20 min, 4°C), the organoids were cryoprotected. The organoids were immersed in 15% (w/v) sucrose in 0.1 M phosphate buffer containing 0.1 M dibasic sodium phosphate and 0.01 M monobasic sodium phosphate in MilliQ water (24 h, 4°C, rotating), and subsequently in 30% (w/v) sucrose in 0.1 M phosphate buffer (48 h, 4°C, rotating). After cryoprotection, the organoids were embedded in freezing medium (15% (w/v) sucrose and 7.5% (w/v) gelatin in 1 M phosphate buffer) and hardened on ice. Freezing was performed in an isopentane bath in liquid nitrogen and the blocks were stored at -30°C until cryosectioning at -18°C into 12 µm–thick sections. The sections were stored at -80°C until use.

### 2.4 Oxygen measurement

The oxygen concentration within the organoids was measured using an optical oxygen microsensor (PM-PSt7, Presens in Regensburg, Germany). The probe was calibrated according to the manufacturer’s instructions. Briefly, the pressure settings were set to the elevation of the lab (997 kPa) and the probe was calibrated in an oxygen-depleted solution (water containing 70 mM sodium sulfite and 500 mM cobalt nitrate) followed by an oxygen-enriched solution (water connected to a room air valve). All solutions were equilibrated to room temperature before calibration and room temperature (20°C) was put as standard in the software. At day 7+18 and day 7+25, organoids in both normoxia and hypoxia were measured using the calibrated sensor. The sensor was fixed to a micromanipulator and was inserted into the bottom of the organoid, approximately 1 mm in depth, close to the insert membrane of the transwell. The incubator was kept closed to ensure a stable gas concentration and measurements were continued until the signal reached a steady state (taking approximately 30–90 minutes).

### 2.5 Hypoxia imaging

Hypoxia was measured in living organoids using the Image-iT Green Hypoxia Reagent (Thermo Fisher Scientific), which produces a green fluorescent signal below 5% O_2_. The organoid was fully immersed in 5 µM ImageIT Green reagent dissolved in organoid culture medium for 4 h in normoxia. Subsequently, the staining solution was replaced with fresh medium and the organoids were moved to the incubator set to either the 7% O_2_ or the 21% O_2_ for 6 h, after which they were fixed for cryosectioning or imaged live. For live imaging, the organoids were imaged using a 10× air objective on an automated Nikon Eclipse Ti-E equipped with a spinning disk with a 70 µm pore size, a Lumencor Spectra X light source and Photometrics Prime 95B sCMOS camera. For cryosectioning, the method in Section 2.3 was applied. The sections were imaged with the same microscope using a 20× air objective and a 40× oil objective. All images were processed using Fiji^34^, in which the rolling ball function was applied in the case of poor signal to noise ratio.

### 2.6 Immunofluorescence

The cryosections were warmed to RT (room temperature) and subsequently incubated in pre-warmed PBS (15 min, 37°C), to remove the sucrose and gelatin. Next, the cryosections were blocked with PBS containing 0.2% (v/v) Tween, 10% (w/v) BSA (bovine serum albumin) and 0.1 M glycine (20 min, RT) and incubated in primary antibodies (Table 1) diluted in the dilution buffer of PBS containing 0.2% (v/v) Tween, 1% (w/v) BSA and 0.1 M glycine (overnight, 4°C). After washing in PBS containing 0.2% (v/v) Tween, the slides were incubated with appropriate secondary antibodies (Table 1) diluted in the dilution buffer (1 h, RT). Finally, the slides were washed and mounted with Prolong Gold (Thermo Fisher Scientific). After curing for two days, imaging was performed on the automated Nikon Eclipse Ti-E microscope using a 20× air and a 40× oil objective. All images were processed using Fiji^34^, in which the rolling ball function was applied in the case of poor signal to noise ratio.

**Table 1:** List of antibodies and stains used for immunofluorescence of cryosections and whole organoids.

| Target | Host | Dilution | Supplier | Cat. no. |
| --- | --- | --- | --- | --- |
| **Primary antibodies and stains** |  |  |  |  |
| CD31 | Sheep | 1:300 | R&D Systems | AF809 |
| E-cadherin (ECAD) | Mouse | 1:300 | BD Bioscience | 610182 |
| HIF1α | Mouse | 1:300 | Abcam | ab16066 |
| HIF2α | Mouse | 1:250 | Santa Cruz Biotechnology | sc-46691 |
| Lotus tetragonolobus lectin (LTL), Biotinylated | N/A | 1:300 | Vector Laboratories | B-1325-2 |
| MEIS 1/2/3 | Mouse | 1:300 | Santa Cruz Biotechnology | SC-101850 |
| Nephrin (NPHS1) | Sheep | 1:300 | R&D Systems | AF4269 |
| SLC21A1 | Rabbit | 1:200 | Sigma-Aldrich | HPAG14967 |
| VEGF-A | Mouse | 1:250 | Santa Cruz Biotechnology | SC-7269 |
| VEGF-A165b | Mouse | 1:500 | R&D Systems | MAB3045 |
| WT1 | Rabbit | 1:500 | Abcam | ab89901 |
| **Secondary antibodies and stains** |  |  |  |  |
| Anti-Mouse 488 | Goat | 1:400 | Thermo Fisher Scientific | 10696113 |
| Anti-Mouse 568 | Goat | 1:400 | Thermo Fisher Scientific | A-11031 |
| Anti-Mouse 647 | Goat | 1:400 | Thermo Fisher Scientific | A-21236 |
| Anti-Rabbit 568 | Donkey | 1:400 | Thermo Fisher Scientific | 10617183 |
| Anti-Sheep 488 | Donkey | 1:400 | Thermo Fisher Scientific | 10473982 |
| Anti-Sheep 568 | Donkey | 1:400 | Thermo Fisher Scientific | A-21099 |
| Goat Anti-Mouse IgG (H + L)-HRP Conjugate | Mouse | 1:3000 | Bio-Rad | 1706516 |
| Streptavidin 647 | N/A | 1:400 | Thermo Fisher Scientific | S21374 |

### 2.7 Luminex assay

The VEGF-A concentration in the culture medium was analyzed using a VEGF-A Human ProcartaPlex Simplex Kit (Cat. no. EPX01A-10277-901, Invitrogen), specific for the detection of VEGF-A165. The medium was collected on days 7+7, 7+12, 7+17, 7+21, and 7+24 and centrifuged (10 min, 4°C, 239 x g). The supernatant was immediately stored at -80°C. The assay was performed according to the manufacturer’s instructions. In brief, samples were diluted 1:50 in universal assay buffer and 50 µL was added to the wells containing the antibody-coupled beads, along with the standards provided with the assay (30 min, RT, shaking). After an overnight incubation (4°C) and a final incubation (30 min, RT, shaking), detection antibody– biotin reporters were added to each well (30 min, RT, shaking). Next, fluorescent conjugate streptavidin–phycoerythrin was added (30 min, RT, shaking). After a final washing step, the beads were resuspended in 120 μL reading buffer. Fluorescence intensities were measured using a Luminex100 instrument (Bio-Rad) which was calibrated before each use. Data acquisition was done with the Bio-Plex Manager 6.0 software. The data were normalized to the standard curve dilutions delivered with the kit according to manufacturer’s instructions.

### 2.8 Western blot

After snap freezing in liquid nitrogen, the organoids were resuspended in 70 µL RIPA (radioimmunoprecipitation assay) lysis buffer (Sigma-Aldrich) supplemented with phosphatase inhibitor tablets PhosSTOP (Sigma-Aldrich) and protease inhibitor cOmplete ULTRA Tablets (Sigma-Aldrich). The protein concentrations were determined using a Pierce BCA Protein Assay Kit (Thermo Fisher Scientific) according to the manufacturer’s protocol. Per well, 15 µg of protein was loaded. The migration was performed in migration buffer (Tris (tris(hydroxymethyl)aminomethane)-EDTA (ethylenediaminetetraacetic acid), SDS (sodium dodecyl sulfate), glycine, Bio-Rad) at 120 V. Subsequently, the proteins were transferred onto a nitrocellulose membrane at 350 mA for 90 minutes in a transfer buffer (Tris-base, glycine, SDS, 20% methanol). After the transfer, the membranes were blocked in blocking buffer (TBS (Tris-buffered saline), 0.1% Tween (v/v), 5% BSA (w/v)) (1h, RT, shaking). Next, the membranes were incubated with primary antibodies (Table 1) (overnight, 4°C, shaking) in TBS-Tween, 5% BSA (w/v). The membranes were washed in TBS-Tween and incubated with the peroxidase-conjugated secondary antibody (Bio-Rad, Table 1) for 1 hour at RT. The membranes were incubated with Clarity Western ECL substrate (Bio-Rad) and were developed using a ChemiDoc (Bio-Rad). The protein bands were quantified by densitometry using ImageJ, normalized to GAPDH.^35^

### 2.9 RNA isolation and qPCR

Organoids stored in TRIzol at -80°C were thawed and pipetted vigorously to homogenize the samples. Then, 500 µL was transferred to a Phasemaker tube (Thermo Fisher Scientific) after which, 100 µL of chloroform were added, shaken thoroughly and incubated for 5 min at RT. The mixture was centrifuged at 12,000 × g (15 min, 4°C). The aqueous phase was carefully transferred to a new microcentrifuge tube containing 250 µL isopropanol and 1 µL glycogen. After an additional centrifugation step (15 min, 4°C, 12,000 × g) the pellet was washed twice with 200 mM NaOAc in 75% ethanol. The supernatant was discarded, and the pellet was dried at 55°C, resuspended in 25 µL nuclease-free water, and incubated at 55°C with shaking. CDNA was synthesized using the iScript cDNA synthesis kit (Bio-Rad) and 500 ng/µL of RNA was loaded per sample. Quantitative PCR was carried out with iQ SYBR Green Supermix (Bio-Rad), 5 ng cDNA per reaction, on a CFX96TM Real-Time system (Bio-Rad). Primer sequences (Table 2) were verified using total human kidney RNA (Takara Bio). *PSMB4* was determined as the most stable housekeeping gene using the method described by Xie, et al. ^36^, and *GAPDH* was used as a second housekeeping gene to ensure valid results. The data were normalized to *PSMB4* and plotted relative to the expression of undifferentiated iPSCs. Each data point represents one organoid and the three data points were from three distinct iPSC differentiations.

**Table 2:**
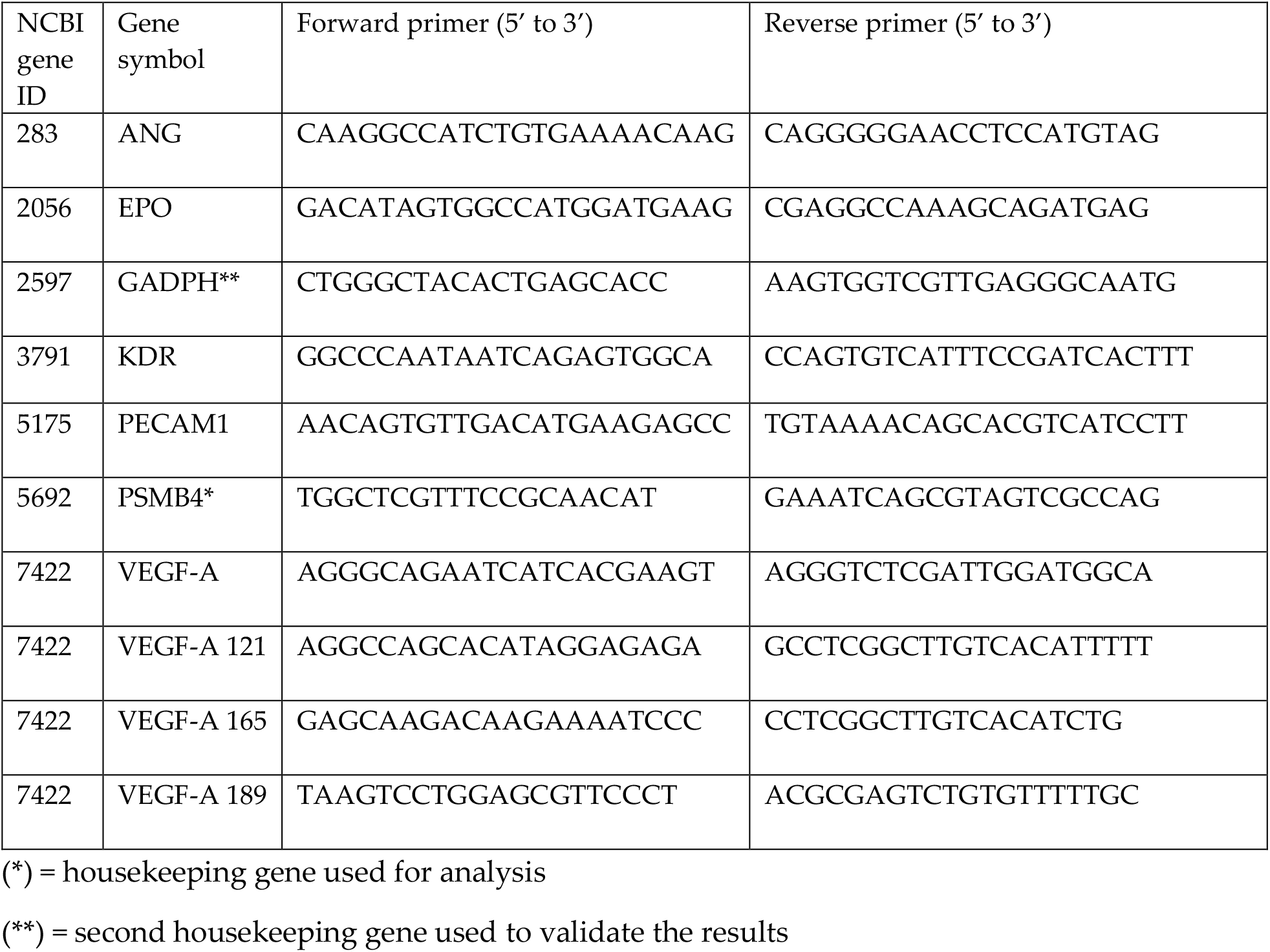
List of qPCR primers

### 2.10 Whole mount immunofluorescence, tissue clearing and automated imaging

Whole organoids were fixed in 2% paraformaldehyde (20 min, 4°C) and blocked in PBS containing 10% goat serum, 0.1 M glycine and 0.5% Triton X-100 (overnight, 4°C, shaking_)_. They were incubated in primary antibodies (Table 1) diluted in PBS containing 10% goat serum, 0.1 M glycine and 0.06% Triton X-100 (3 days, 4°C, shaking). After two washes in 0.3% Triton X-100 in PBS (2 h, RT, shaking), the organoids were incubated with the appropriate secondary antibodies (Table 1) diluted in PBS containing 1% goat serum, 0.1 M glycine and 0.06% Triton X-100 (3 days, 4°C, shaking). After two washes (2 h, RT, shaking), the organoids were cleared using the method of Klingberg, et al. ^37^. Briefly, the organoids were dehydrated in a series of 30% to 100% molecular biology–grade ethanol and submerged in ethyl cinnamate (overnight, RT followed by 1 h, 37°C). Imaging was done immediately after on an automated Nikon Eclipse Ti-E with the 70 µm spinning disk in place. Imaging was automated using a Nikon JOBS tool. Briefly, a 10× air objective was used for automated detection of the organoid, while a 20× air objective with an extra-long working distance was used to image at higher resolution with a z increment size of 5 µm.

### 2.11 Automated image processing and vessel quantification

Whole mount images were processed within a processing and quantification JOB specifically made for these samples, integrated in the NIS-Elements AR (Advance Research) (Nikon) software. In summary, background was removed using the clarify AI tool prior to the segmentation of the CD31 signal. The CD31 signal was thresholded by intensity and segmented by measuring the longest medial axis in 3D. Falsely segmented pixels, due to intense background signal, were excluded by circularity and minimal size if needed. The volume of the segmented signal was measured in 3D for the full organoid. The volume of the total organoid was measured by segmenting the organoid based on the strong background. The percentage of vascularization was determined by calculating the CD31 volume as a fraction of the total organoid volume. Furthermore, the segmented CD31 signal was skeletonized to measure the cumulative length of all fragments in 3D. Stepwise details on the used JOB can be found in Supplementary Figure 1.

### 2.12 Statistical analysis

All statistical analyses were performed using GraphPad Prism 9. For all immunofluorescence protein analyses, three organoids (n=3) each arising one of three independent organoid cultures (N=3) were assessed. For the gene expression analysis, Western blot and Luminex assay, three organoids (n=3) each arising from one of two independent organoid cultures (N=2) were analyzed. Two organoids (n=2), each arising from one of three independent organoid cultures (N=3) were assessed for the darkfield measurement and two organoids (n=2) each arising from one of two independent organoid cultures (N=2) were used for the oxygen measurement. Individual samples were excluded only for technical failures. For each figure, the exact N and n are reported in the figure caption. A two-way ANOVA was performed for each experiment to assess the contribution of both row (time point) and column (oxygen concentration) factors. All *p*-values can be found in the Results section. Statistical significance was only concluded for *p*-values below 0.05.

## 3 Results

### Macro-morphology and oxygen concentrations in normoxic and hypoxic organoid culture

To replicate the *in vivo* hypoxic environment, we cultured kidney organoids in 7% O_2_ and compared them to their counterparts cultured in 21% O_2_ (Figure 1A). Brightfield analysis revealed no differences in macro morphology, density or size until day 7+25 (Figure 1B, Supplementary Figure 2). Darkfield imaging was performed to assess the shape and thickness of the organoids (Figure 1C). No statistically significant differences were found comparing organoids cultured in normoxia and hypoxia (*p* = 0.174; Figure 1D).

To quantify the oxygen concentration, we inserted an optical microsensor into the bottom of the organoid (close to the filter). In normoxia, the lowest oxygen concentration we measured was 5.83 ± 1.03% on day 7+18, which significantly decreased to 1.55 ± 0.40% by day 7+25 (*p* = 0.002). This value at day 7+25 was not significantly different from the oxygen concentration measured in the organoids cultured in hypoxia on day 7+18 (1.33 ± 0.80%; *p* = 0.983) or day 7+25 (1.00 ± 0.27%; *p =* 0.837) (Figure 1E).

To investigate whether the organoids had a hypoxic core, we imaged living organoids and cryosections at the center of the organoids with a hypoxic dye (ImageIT Green) (Figure 1F). The cryosections, cut through the center of the organoids, revealed a higher intensity (meaning lower O_2_) in hypoxia. We saw the dye localized largely to nephron structures and less to the surrounding stroma. We did not see evidence of a hypoxic core in the center of any organoids.

Next, we explored if culture in hypoxia would affect the differentiation and organization of various cell types in the kidney organoids. To assess this, we stained vertically cut cryosections throughout the whole organoids for well-established markers to label the major cell types found in kidney organoids. We found no differences in either the presence of the cell types or their organization into proximal tubules (LTL), distal tubules (ECAD), loop of Henle (SLC12A1) or glomeruli (NPHS1-positive podocytes) (Figure 2A–B).

**Figure 2.**
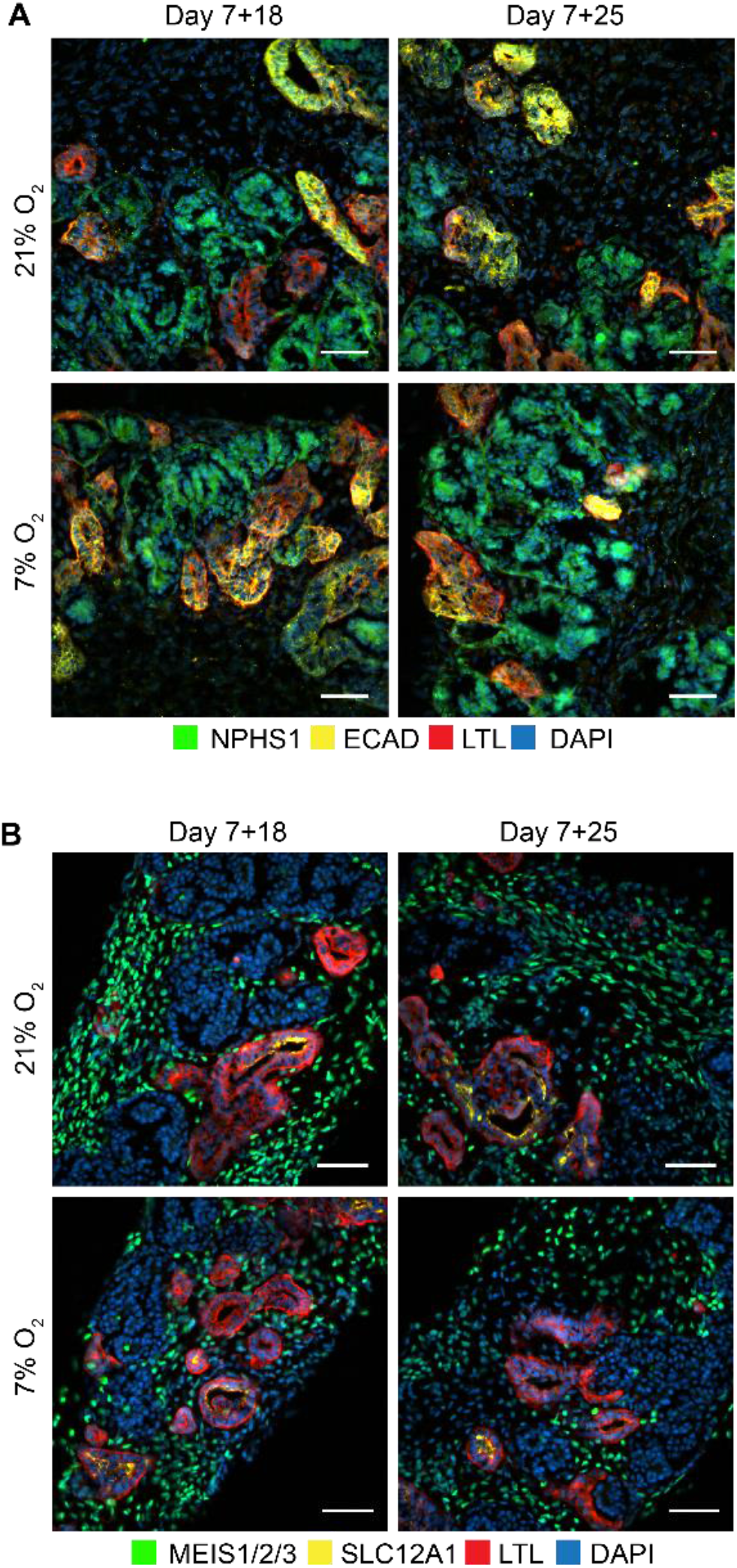
Kidney organoids cultured at 7% and 21% O_2_ develop the same cell types. Immunofluorescence staining for A. podocytes (NPHS1; green), proximal tubules (LTL; red) and distal tubules (ECAD; yellow) and nuclei (DAPI; blue), and B. interstitial cells (MEIS1/2/3; green), loop of Henle (SLC12A1; yellow) and proximal tubules (LTL; red), showed these different cell types in all conditions. (n=3, N=3). Scale bars: 50 µm.

### Nephrons express varied levels of nuclear HIF1α and a constant nuclear expression of HIF2α

Nuclear HIF translocation during kidney development is crucial for nephrogenesis and angiogenesis. We investigated which cells showed responsiveness to hypoxia in terms of nuclear translocation of HIF1α and HIF2α. For this, we performed immunofluorescence staining of HIF1α and HIF2α on vertically cut cryosections throughout the whole kidney organoids at days 7+18 and 7+25. Qualitative analysis showed that HIF1α was differentially expressed in hypoxia and normoxia. At day 7+25, organoids in the hypoxia culture had increased HIF1α nuclear expression compared to those in normoxia (Figure 3A). Particularly, the podocytes had more nuclear translocation in both normoxia and hypoxia compared to other cell types. Furthermore, in normoxia, podocytes at the bottom of the organoid, towards the medium interface, expressed nuclear HIF1α, while in normoxia this was seen in podocytes throughout the whole organoid. In all conditions and time points, interstitial cells and most tubules did not express nuclear HIF1α. HIF2α was expressed in the nuclei of all cells of the kidney organoids at days 7+18 and 7+25 (Figure 3B). Particularly the interstitial cells had heterogeneous expression. We did not detect differences between organoids cultured in normoxia or hypoxia in terms of HIF2α nuclear translocation.

**Figure 3.**
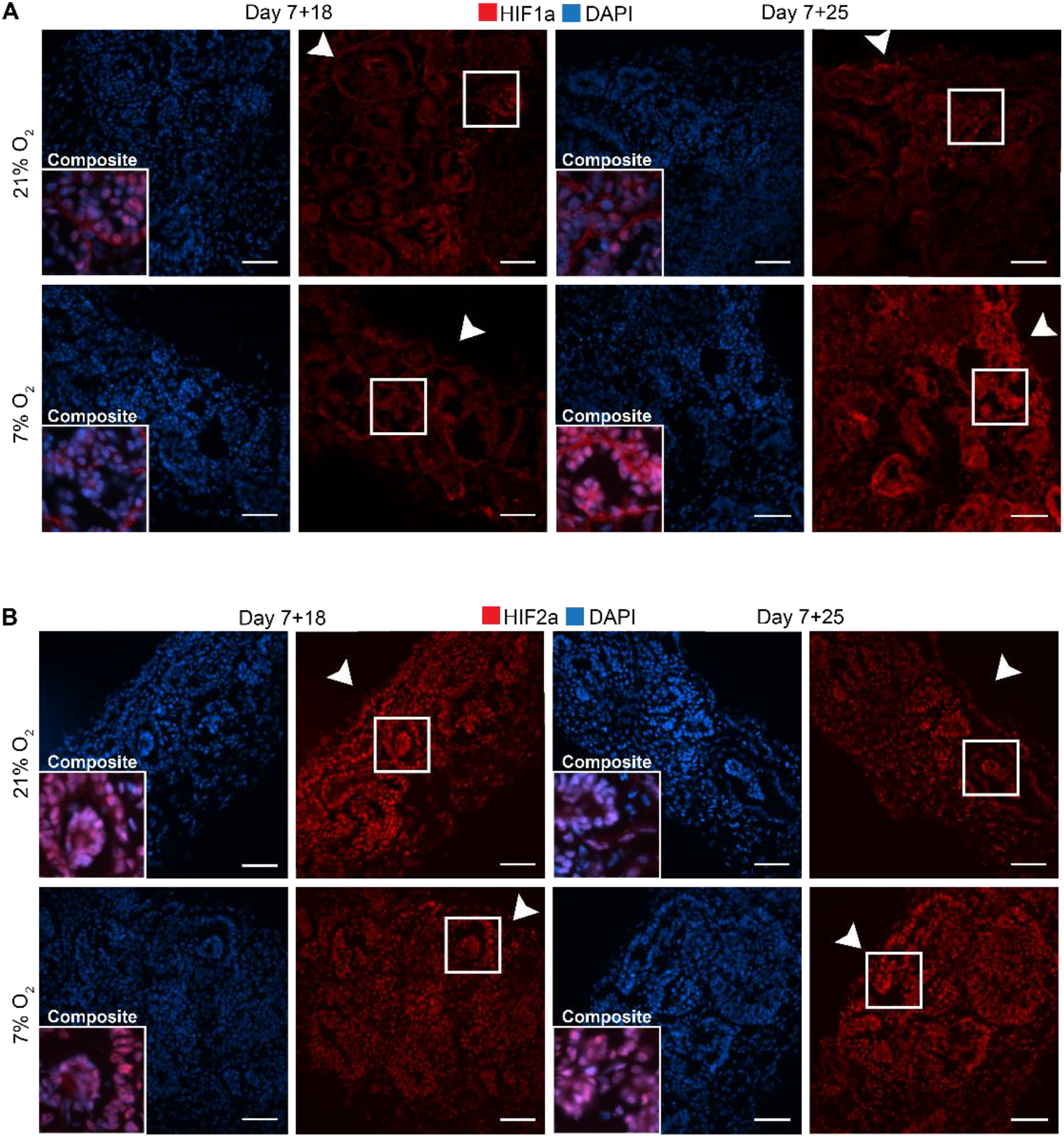
Kidney organoids cultured in both 7% and 21% O_2_ show differential nuclear translocation of HIF1a and HIF2a. A. Immunofluorescence staining for HIF1a (red) and DAPI (blue) shows increased nuclear translocation in podocytes (composite), particularly in 7% O_2_ at day 7+25. Scale bar: 50 µm. B. HIF2a (red) and DAPI (blue) immunofluorescence staining shows nuclear translocation in most cells within the organoid irrespective of the oxygen concentration or time point. (n=3, N=3). Scale bar: 50 µm. Arrowheads indicate the (air-exposed) surface of the organoids.

### VEGF-A protein expression in proximal tubules

VEGF-A is a major angiogenic factor primarily produced in distal tubules, collecting duct and podocytes.^38^ It is regulated by hypoxia by the binding of nuclear, dimerized HIFs to the HRE promotor. We performed immunofluorescence on vertically cut cryosections throughout whole organoids to determine if there was a difference in expression between normoxia- and hypoxia-cultured organoids (Figure 4A), and which cell types expressed VEGF-A (Figure 4B). We found expression on the apical side of some tubular structures, mainly in normoxia and to a lesser extent in hypoxia (Figure 4A-B), but no difference between day 7+18 and day 7+25. The VEGF-A+ tubules in both normoxia and hypoxia co-expressed both SLC12A1 marking the loop of Henle (thick ascending limb) and LTL marking proximal tubules (Figure 4B). We validated this co-expression of SLC12A1, LTL and VEGF-A on adult kidney sections and in scRNA sequencing datasets of organoids (Supplementary Figure 3). Glomerular expression of VEGF-A in the normoxia and hypoxia organoids was not found. We hypothesized that the lower VEGF-A expression in the proximal tubules in hypoxia could be due to a switch to a more soluble VEGF-A isoform or that the time point of secretion would be different. Therefore, soluble VEGF-A was measured using a Luminex assay validated for the VEGF-A165 isoform in the culture medium on days 7+7, 7+12, 7+17, 7+21, 7+24 (Figure 4C). We found a significant increase over time from day 7+7 until 7+17 (*p* ≤ 0.0002) for organoids in both hypoxia and normoxia. After day 7+17, the soluble VEGF-A decreased significantly until day 7+21 (*p* ≤ 0.012). There was no significant difference between organoids in normoxia or hypoxia (*p* = 0.544).

**Figure 4.**
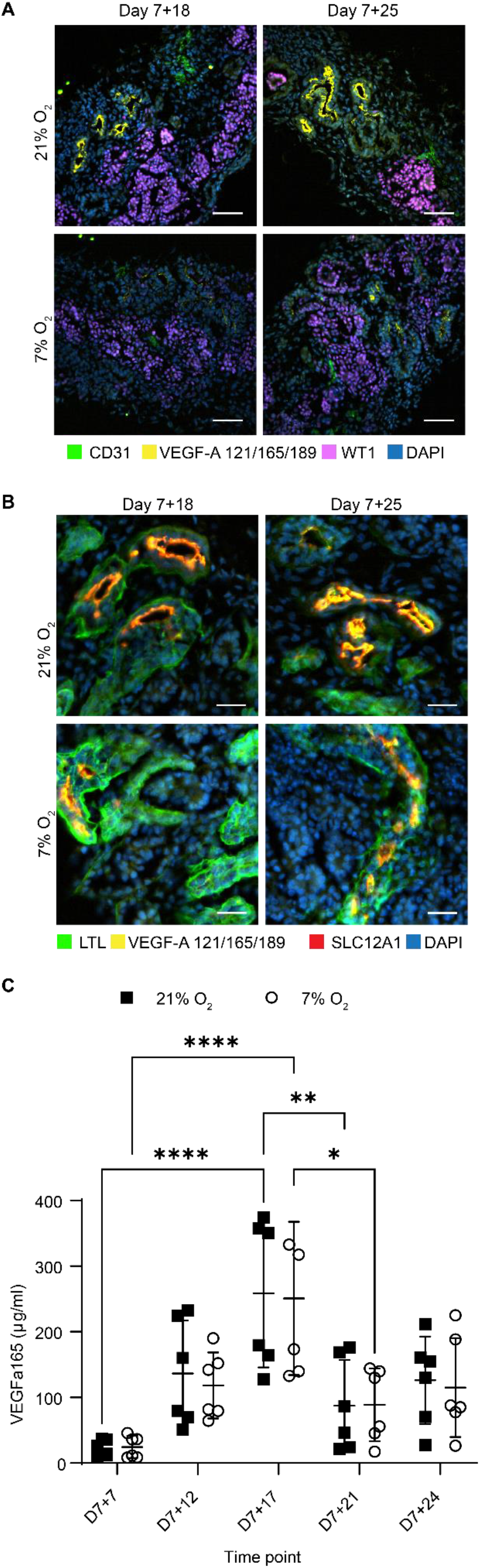
Expression of hypoxia-responsive VEGF-A in organoids cultured in 21% and 7% O_2_. A. Immunofluorescence for VEGF-A (121, 165, 189 isoforms; yellow), endothelial cells (CD31; green), podocytes (WT1; magenta) and nuclei (DAPI; blue) displays a reduction of VEGF-A in 7% O_2_ compared to 21% O_2_ at both day 7+18 and day 7+25. (n=3, N=3). Scale bar: 50 µm. B. Immunofluorescence shows localization of VEGF-A to the apical side of proximal tubules co-positive for loop of Henle marker (SLC12A1; red). (n=3, N=3). Scale bar: 25 µm. C. VEGF-A165 protein expression measured in the culture medium was differentially expressed over time, but not significantly different in hypoxia compared to normoxia. (n=3, N=2). *= *p* ≤ 0.05, ** = *p* ≤ 0.007, **** = *p* < 0.0001.

### Differential expression of angiogenesis-regulating genes in hypoxic culture

*VEGF-A* is known to exist in various splice variants with distinct biological functions. *VEGF-A165* is the most prominent variant, followed by *189* and *121* in the adult human kidney.^39^ We investigated *VEGF-A* variant expression in the organoids by quantitative polymerase chain reaction (qPCR). *VEGF-A165* was significantly upregulated from day 7+18 to day 7+25 (*p* = 0.049), but there was no significant difference between organoids in normoxia or hypoxia (Figure 5B). *VEGF-A189* was also significantly upregulated in hypoxia over time (Figure 5C). Furthermore, *VEGF-A189* was significantly higher in hypoxia cultures than in normoxia, with a 2.5 fold-change (FC) and 1.5 FC, respectively, at day 7+18 relative to expression in control iPSC cultures (*p* < 0.001; Figure 5C). This difference in VEGF-A189 upregulation was also observed at the day 7+25 timepoint between hypoxia and normoxia cultures, with a 3.5 FC and 2.7 FC respectively (*p* = 0.021), as plotted relative to the control iPSCs. For *VEGF-A121*, we found no difference in its expression between normoxia or hypoxia cultures or over time (Figure 5D).

**Figure 5.**
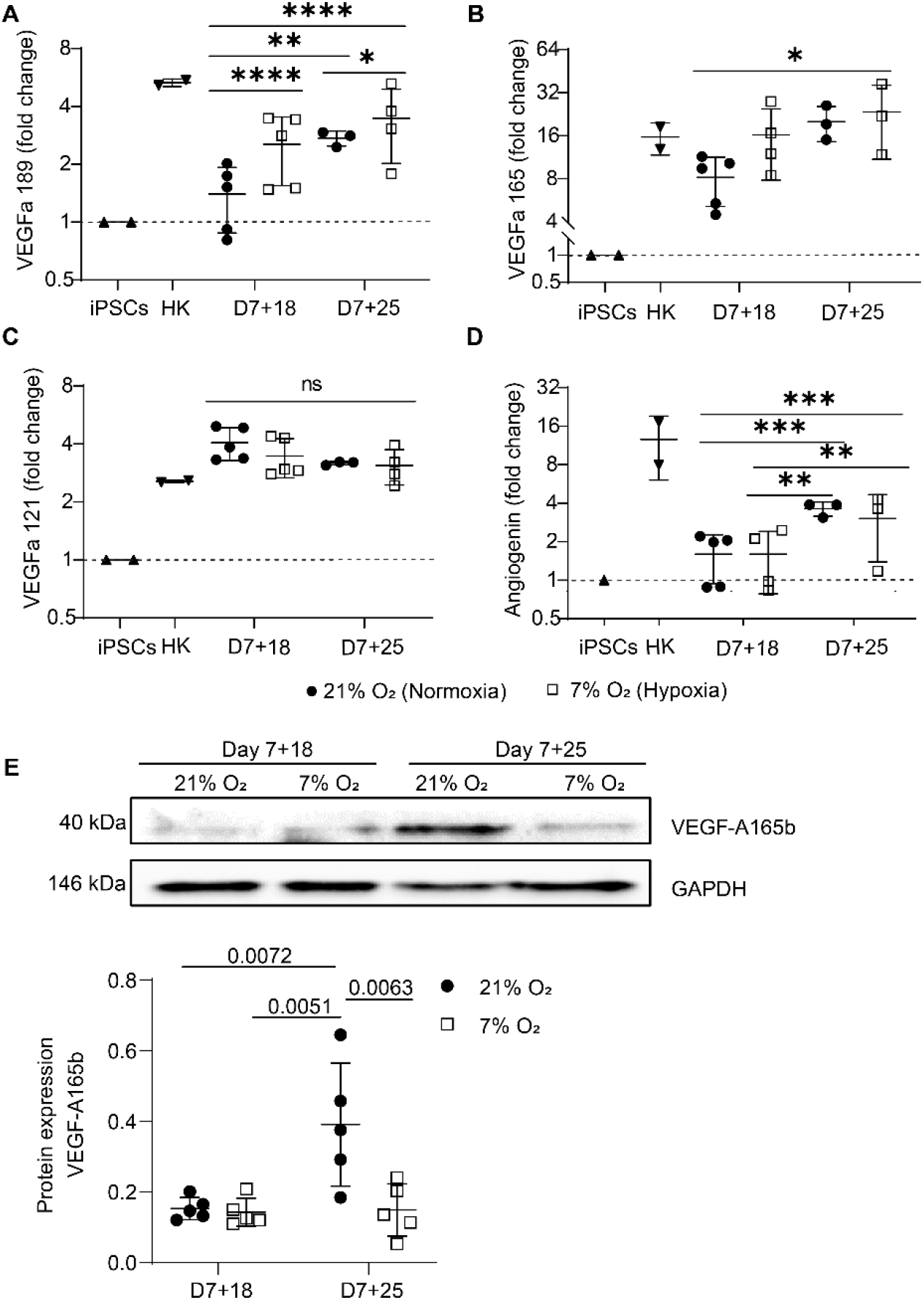
Hypoxic organoid culture differentially upregulates VEGF-A variants. A. *VEGF-A189* and B. *VEGF-A165* are upregulated in 7% O_2_, while C. *VEGF-A121* remains unchanged. D. Angiogenin mRNA is significantly upregulated over time in both the normoxic and hypoxic culture. For panels A–D, data are normalized to expression in control iPSCs. Total human kidney (HK) RNA is plotted as a reference. (n=3, N=2). E. The anti-angiogenic VEGF-A165b isoform is significantly upregulated over time in normoxia and significantly downregulated in hypoxia compared to normoxia at day 7+25. (n=3, N=2). **= *p* ≤ 0.01, *** = *p* ≤ 0.001, **** = *p* ≤ 0.0001.

We also determined mRNA expression of *ANG*, the gene encoding angiogenin, another hypoxia-regulated angiogenic factor, in our kidney organoid culture. Similar to *VEGF-A165, ANG* mRNA was significantly upregulated over time (*p* < 0.0001) but with no differences between hypoxia and normoxia cultures (Figure 5E). The protein expression of the antiangiogenic isoform VEGF-A165b normalized to GAPDH expression was determined by Western blot (Figure 5E). VEGF-A165b was significantly upregulated in normoxia from day 7+18 to day 7+25 (*p* = 0.007), while there was a significant reduction in hypoxia compared to normoxia at day 7+25 (*p* = 0.006).

### Hypoxia-induced angiogenesis: homogenous micro-vessel formation and sprouting

Hypoxia is well known to enhance angiogenesis in both development and pathologies.^40^ In both, endothelial cells undergo sprouting to form new blood vessels. We therefore investigated the effect of hypoxia on the endothelial patterning by whole mount immunostaining with the endothelial marker CD31. Whole mount imaging revealed enhanced micro-vessel formation with more homogenous morphology, enhanced branching and sprouting in organoids cultured in hypoxia compared to normoxia (Figure 6A, C). We observed a reduced intensity of CD31 (not quantified) in hypoxia and normoxia at day 7+25 (Figure 6B). However, the endothelial phenotype was maintained, as confirmed by co-expression of CD31 and VE-cadherin in both normoxia- and hypoxia-cultured organoids at day 7+25 (Supplementary Figure 4). We set up an automatic 3D segmentation and quantification pipeline and found that hypoxia induced a significant increase in the fraction of endothelial cells (40.3 ± 12.50%) compared to normoxia (14.3 ± 2.60%) (*p* = 0.032; Figure 6E) at day 7+18 that was not observed at day 7+25 (*p* = 0.226). Interestingly, the average vessel length was significantly higher in hypoxia compared to normoxia at day 7+18 (hypoxia: 158 ± 12 µm; normoxia: 105 ± 18 µm; *p* = 0.028) and day 7+25 (hypoxia: 154 ± 13 µm; normoxia: 97 ± 11 µm; *p* = 0.036) (Figure 6D). Three-dimensional segmentation additionally revealed that the endothelial network in organoids cultured in normoxia largely resided in a two-dimensional plane (parallel to the transwell), while in hypoxia culture, the network was mainly interconnected in three dimensions (supplementary movies).

**Figure 6:**
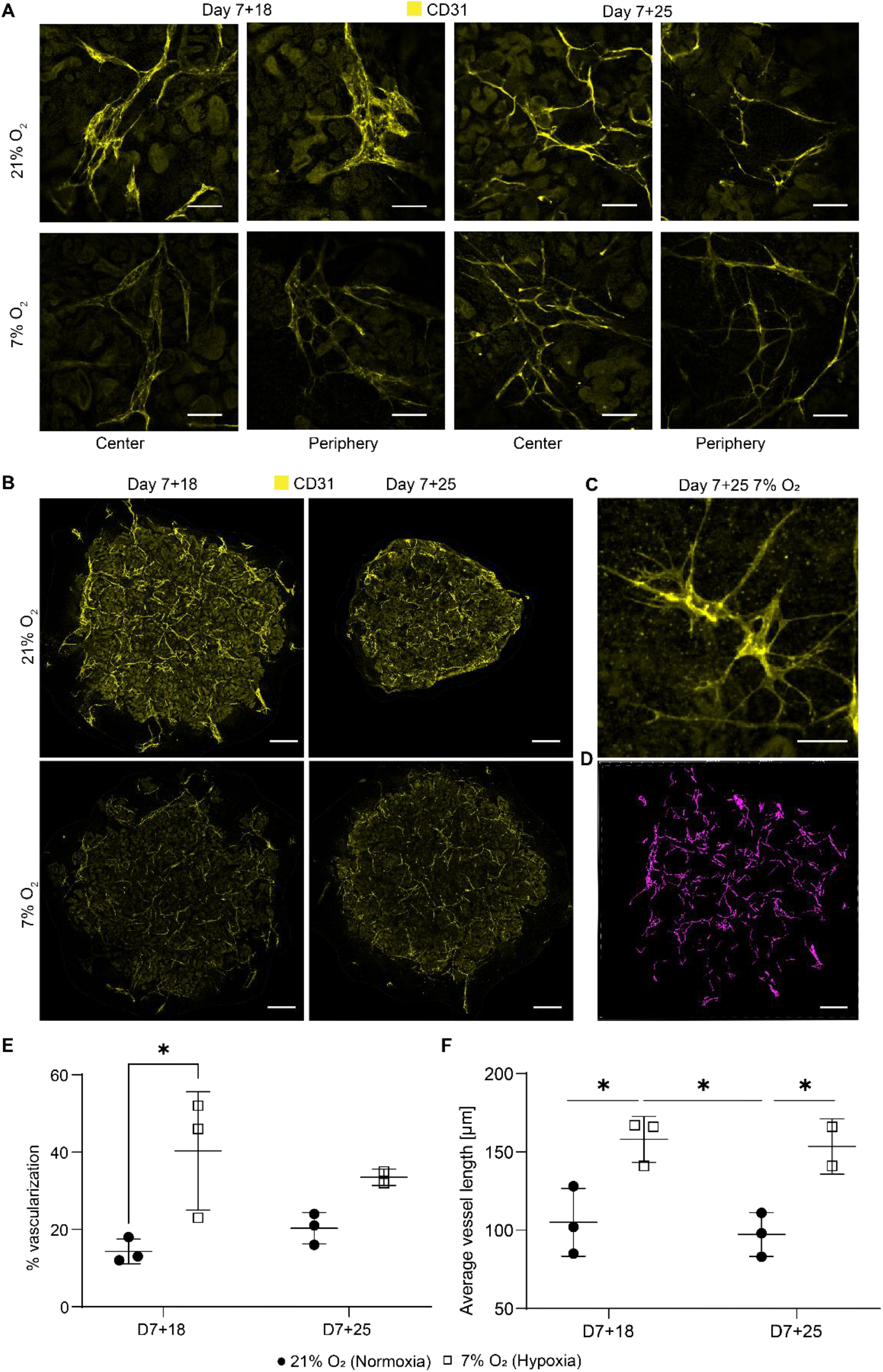
Hypoxic organoid culture promotes interconnected microvasculature formation and sprouting. A. The endothelial network (CD31; yellow) has a more homogeneous morphology and enhanced interconnectivity and branching in organoids cultured in 7% O_2_. B. Whole mount imaged organoids show less intense CD31 staining, likely due to smaller vessel size. C. Hypoxia-induced sprouting of endothelial vessels at day 7+25. D. Example of 3D segmentation of endothelial network of a day 7+18 normoxia organoid. Scale bars: 50 µm (A, C); 600 µm (B, D). Quantification in 3D reveals E. an increase in the volume fraction of endothelial cells at day 7+18 in hypoxia and F. an increased average vessel length in hypoxia compared to normoxia. (n=1, N=3). * = *p* ≤ 0.036.

## 4 Discussion

Our aim was to investigate if a lower oxygen concentration applied to the kidney organoid culture that mimics the *in vivo* hypoxic environment during nephrogenesis could improve angiogenesis and organoid vascularization. The mRNA expression of endothelial markers (*CD31, KDR*) in the normoxic culture was downregulated over time (Supplementary Figure 5), indicating the need for stimuli to activate angiogenesis. We opted for a long-term hypoxic culture (20 days), because long-term hypoxia (hours to days, depending on the model) enhances the expression of angiogenic cytokines and growth factors such as PDGF and VEGF in comparison to acute hypoxia that is known to activate the release of inflammatory factors.^41,42^ VEGF-A in particular is known to be crucial for endothelial cell differentiation, proliferation and migration^43^, as well as glomerular capillary formation, podocyte survival and slit diaphragm integrity in autocrine podocyte signaling.^44^ We therefore included VEGF-A mRNA and protein expression in our investigations.

Kidney organoids developed similarly in normoxic and hypoxic conditions. The hypoxic culture did not impair nephron specification and organization, nor organoid size (Figure 1B– C, Figure 2A–B), indicating that hypoxia did not interfere with normal organoid development. Our measurements of oxygen concentration on the bottom of the organoid culture plate (1.0 ± 0.3% in hypoxia on day 7+25) were lower compared to the poorly vascularized medullas in adults (1.9% O_2_ at atmospheric pressure derived from the measured 15 mmHg).^28^ We therefore expected our hypoxic culture to be comparable to avascular, hypoxic kidneys during early development.^45^ This similarity indicates that growing organoids at 7% O_2_ could mimic the *in vivo* hypoxic environment and help study organoid maturation in a more physiological model.

In addition to oxygen levels, we found a similar transcriptional program activated in the kidney organoids as in the developing human kidney. Specifically, severe hypoxia and consequently HIF stabilization was found to be essential in nephrogenesis^38^, with HIF-1α expressed in cortical and medullary collecting duct, nephrogenic zone and glomerular cells.^30^ While the collecting duct lineage does not exist in the organoids, podocytes deeper within the organoid, showed nuclear HIF-1α expression at day 7+18 in normoxia (Figure 3A). According to our measurements, these podocytes would reside in regions close to 5.83 ± 1.03% O_2_. Likely, only podocytes experiencing less than 5% O_2_ showed nuclear translocation of HIF-1α.^27^ Indeed, even peripheral podocytes of the hypoxia-cultured organoids, where the oxygen concentration was 1.0 ± 0.27% at the bottom of the organoids, showed nuclear HIF-1α translocation (Figure 3A). While certainly not all podocytes responded by nuclear HIF-1α translocation, the hypoxic culture clearly induces nuclear HIF-1α in podocytes throughout the organoids.

Comparable to HIF-1α, nuclear HIF-2α is important to *in vivo* kidney development, where it is known to be expressed only in interstitial cells and podocytes^13^, as shown in week 14 fetal kidneys^30^, as well as in developing tubules in newborns^38^. In our kidney organoids, HIF-2α expression was not limited to interstitial cells and podocytes, but was expressed in all cell types examined in both hypoxia and normoxia (Figure 3B). There were clear differences in intensity of nuclear HIF-2α in interstitial cells within the organoids in normoxia and hypoxia, while the nephrons equally expressed nuclear HIF-2α. The reason for this observed difference is unclear. Clarification is also needed for why all interstitial cells, even in the most oxygenated condition (day 7+18 normoxia), do express nuclear HIF-2α. There is a need for further investigation, since recent findings in mice suggest that chronic HIF-2α expression in stromal progenitors impairs kidney development, in particular nephron formation, tubular maturation, and the differentiation of FOXD1+ stromal cells into smooth muscle, renin, and mesangial cells.^46^ This was found to be regulated by the inhibition of PhD2 and PhD3.^47^ The fact that the organoids do not mature further and mesangial cells and renin cells are thought to be absent in this kidney organoid model^48^, could indicate impaired *in vitro* differentiation of FOXD1+ progenitor cells. Therefore, future research could investigate the role of HIF-2α nuclear translocation in this context.

*In vivo*, pericytes derived from FOXD1+ progenitors are known to show HIF-2α nuclear translocation that induces EPO production. However, in our organoids, HIF-2α nuclear translocation did not induce EPO mRNA transcription in either normoxia or hypoxia (Supplementary Figure 6). The fact that EPO was not transcribed in the organoids in normoxia and hypoxia could potentially be due to a fibrotic stromal population found in the organoids^33^ and subsequent hyper-methylation of the EPO 5’ and HRE, inhibiting HIF2/HIFβ dimer binding, as found in adult fibrotic kidneys^49^. Future research could clarify mechanistically, if nuclear HIF-2α is actually leading to target gene transcriptions or if this is inhibited, consequently being one reason for limited organoid maturation.^46^

As with HIF-2α expression, we found differences in the expression of VEGF-A in our kidney organoids compared to fetal human developing kidneys. Comparing the normoxic and hypoxic organoid culture, a decrease in VEGF-A protein expression could be seen. VEGF-A, being one of the most important angiogenic factors, is known for its function in nephrogenesis to induce blood vessel formation in glomeruli through branching angiogenesis and consequently to induce maturation of glomerular cells.^50^ In fetal human developing kidneys, VEGF-A is expressed in the epithelial cells in the s-shaped body and collecting duct.^29^ Later in nephrogenesis, VEGF-A uptake by convoluted tubules has been observed.^51^ In the organoids, VEGF-A was localized on the apical side of LTL+ proximal tubules, co-expressing the LoH marker SLC12A1. We did not detect its expression in podocytes, which is needed to initiate the formation of glomerular capillaries.^52^ Consequently, glomeruli containing endothelial cells is a rare event in the organoids. In hypoxia, the signal for VEGF-A was less intense, indicating reduced expression or absorption by the proximal tubule (Figure 4 A–B). We hypothesized that there could be differences in the VEGF-A isoform and transcript variant expression, which could remain hidden by targeting three isoforms with the same antibody as performed in Figure 4A and B. *In vivo*, podocytes are the main source of VEGF-A and synthesize three VEGF-A isoforms (VEGF-A-121, VEGF-A-165, VEGF-A-189) by alternative mRNA splicing.^44^

VEGF-A variants were differentially expressed in the organoids in hypoxia (Figure 5A–C), consistent with previous studies.^53^ *VEGF-A189* was significantly upregulated in organoids cultured in hypoxia, which is associated with microvessel formation^39^ and enhanced migration of endothelial cells^54^. Being positively charged in some domains, encoded for by exons 6a, 6b and 7, VEGF-A189 binds negatively charged extracellular matrix and heparan sulphate proteoglycans on cell surfaces, remaining spatially localized and becomes biologically active upon mobilization by heparinase.^55,56^ This allows the attraction of vessels into hypoxic tissue. Enhanced cell migration is also confirmed in a variety of cell lines cultured at 0.5% O_2_ and modified to overexpress VEGF-A189.^57^ While we could not prove the localization of this isoform in the organoids due to unavailability of isoform-specific antibodies, we did find an improved patterning of the endothelial network in hypoxia compared to normoxia. Microvessels were homogenously sprouting (Figure 6A–C) with larger vessel length (Figure 6F) and larger connectivity in 3D (Supplementary movies) in hypoxia. In normoxia, there was comparatively less connectivity, hetereogenous vessel morphologies and planar growth.

The VEGF-A165b isoform was downregulated in hypoxia compared to normoxia (Figure 5E). VEGF-A165b is a low efficacy agonist, binding VEGFR2 and thereby reducing binding of VEGF-A165, resulting in strongly decreased signal transduction via the VEGFR2 receptor.^58^ VEGF-A165b is upregulated in quiescent vessels, inhibiting endothelial cell migration^51^, and in adult kidneys^58^ and is downregulated in nephrogenesis during capillary loop formation^51,59^. Downregulation of the VEGF-A165b isoform in hypoxia at day 7+25 could indicate increased binding of VEGF-A165 to the VEGFR2 receptor, enhanced signal transduction and consequently the initiation of angiogenesis. Earlier research confirms the phosphorylation of SRSF1 splice factor by hypoxia targeting the exon 8a proximal splice site, leading to the expression of angiogenic VEGF-A isoforms.^53^ While the increased vessel sprouting observed in the hypoxia-cultured organoids is an indication of angiogenesis^60^ (Figure 6A–C), more research is needed to prove causality with the downregulated VEGF-A165b isoform. Furthermore, to our knowledge, alternative splicing of VEGF-A in non-pathological developmental angiogenesis is insufficiently studied and would be highly valuable in the context of organoid vascularization and maturation. Finally, *in vivo* glomerular maturation is VEGF-A dose dependent, however, it is only hypothesized that the antiangiogenic isoforms have a dose dependent effect in glomerulogenesis as well.^51^

The results of our study show that modulation of the oxygen concentrations in kidney organoid culture can improve the patterning of endothelial cells and therefore is potentially a relevant factor to integrate in regular organoid culture. Sprouting and interconnected vessels in hypoxia indicated the activation of angiogenesis, which is important for further nephron maturation. Future research is, however, needed to confirm the mechanism by which this enhanced patterning of the endothelial network takes place and to elucidate the roles of VEGF-A189 and VEGF-A165b. This understanding will help to further enhance the endothelial network in kidney organoids and can potentially be applied to other systems as well. Furthermore, we hypothesize that a culture in a hypoxic chamber could be a more controlled environment to study the effects of a hypoxic culture on organoid development. This setup would allow medium changes at the desired hypoxia instead of at ambient oxygen concentrations and would consequently avoid repeated reoxygenation of the organoids. Finally, it remains to be determined how VEGF-A expression by podocytes can be increased to allow vascularization and maturation of glomeruli.

## 5 Conclusion

In conclusion, kidney organoid culture in physiological hypoxia induced the formation of a homogenous and interconnected endothelial network, while maintaining renal cell types and their spatial organization. We found that *VEGF-A189* mRNA is upregulated in hypoxia, which has been associated with the formation of microvessels in hypoxic tissue in earlier studies. Furthermore, the antiangiogenic isoform VEGF-A165b was significantly downregulated in hypoxia, being a potential reason for the enhanced sprouting. While further vessel maturation, i.e. tube formation and glomerular vascularization, are still unresolved, we believe that culture in physiological hypoxia is an important driving force for organoid vascularization and is translatable to other organoid models in a model-specific manner.

## Acknowledgements

The authors wish to thank the Physiology department at the Maastricht University for the opportunity to use their hypoxia incubator and Hang Nguyen for reviewing this manuscript. Furthermore, the authors would like to thank Carlos Julio Peniche Silva and Daniel Carvalho from the MERLN Institute for initial support with the oxygen measurements, Sven Hildebrand from Cognitive Neuroscience at Maastricht University for vivid discussions on tissue clearing and Jasmin Dehnen for the support with designing variant-specific primer pairs. Figure 1A was created with BioRender.com.

## Disclosure of Potential Conflicts of Interest

The authors declare no competing interests.

## Data Availability Statement

The raw and processed data required to reproduce these findings are available to download from Dataverse.

## Notes

### Competing Interest Statement

The authors have declared no competing interest.

## References

1 Takasato, M., Er, P. X., Chiu, H. S. & Little, M. H. Generation of kidney organoids from human pluripotent stem cells. Nat Protoc 11, 1681–1692, doi:10.1038/nprot.2016.098 (2016).

2 Schumacher, A. et al. Defining the variety of cell types in developing and adult human kidneys by single-cell rna sequencing. NPJ Regen Med 6, 45, doi:10.1038/s41536-021-00156-w (2021).

3 Gupta, N., Dilmen, E. & Morizane, R. 3d kidney organoids for bench-to-bedside translation. J Mol Med (Berl) 99, 477–487, doi:10.1007/s00109-020-01983-y (2021).

4 Nishinakamura, R. Human kidney organoids: Progress and remaining challenges. Nat Rev Nephrol 15, 613–624, doi:10.1038/s41581-019-0176-x (2019).

5 Takasato, M. & Wymeersch, F. J. Challenges to future regenerative applications using kidney organoids. Curr. Opin. Biomed. Eng. 13, 144–151, doi:10.1016/j.cobme.2020.03.003 (2020).

6 Rossi, G., Manfrin, A. & Lutolf, M. P. Progress and potential in organoid research. Nat Rev Genet 19, 671–687, doi:10.1038/s41576-018-0051-9 (2018).

7 Place, T. L., Domann, F. E. & Case, A. J. Limitations of oxygen delivery to cells in culture: An underappreciated problem in basic and translational research. Free Radic Biol Med 113, 311–322, doi:10.1016/j.freeradbiomed.2017.10.003 (2017).

8 Ma, T., Grayson, W. L., Frohlich, M. & Vunjak-Novakovic, G. Hypoxia and stem cell-based engineering of mesenchymal tissues. Biotechnol Prog 25, 32–42, doi:10.1002/btpr.128 (2009).

9 Worsdorfer, P. & Ergun, S. The impact of oxygen availability and multilineage communication on organoid maturation. Antioxid Redox Signal 35, 217–233, doi:10.1089/ars.2020.8195 (2021).

10 Okkelman, I. A., Foley, T., Papkovsky, D. B. & Dmitriev, R. I. Live cell imaging of mouse intestinal organoids reveals heterogeneity in their oxygenation. Biomaterials 146, 86–96, doi:https://doi.org/10.1016/j.biomaterials.2017.08.043(2017).

11 Ding, B. et al. Three-dimensional renal organoids from whole kidney cells: Generation, optimization, and potential application in nephrotoxicology in vitro. Cell Transplant 29, 963689719897066, doi:10.1177/0963689719897066 (2020).

12 Grobstein, C. Trans-filter induction of tubules in mouse metanephrogenic mesenchyme. Exp Cell Res 10, 424–440, doi:10.1016/0014-4827(56)90016-7 (1956).

13 Buchholz, B., Schley, G. & Eckardt, K. U. The impact of hypoxia on nephrogenesis. Curr Opin Nephrol Hypertens 25, 180–186, doi:10.1097/MNH.0000000000000211 (2016).

14 Li, J., Gao, X., Qian, M. & Eaton, J. W. Mitochondrial metabolism underlies hyperoxic cell damage. Free Radic Biol Med 36, 1460–1470, doi:10.1016/j.freeradbiomed.2004.03.005 (2004).

15 Jagannathan, L., Cuddapah, S. & Costa, M. Oxidative stress under ambient and physiological oxygen tension in tissue culture. Curr Pharmacol Rep 2, 64–72, doi:10.1007/s40495-016-0050-5 (2016).

16 Brueckl, C. et al. Hyperoxia-induced reactive oxygen species formation in pulmonary capillary endothelial cells in situ. Am J Respir Cell Mol Biol 34, 453–463, doi:10.1165/rcmb.2005-0223OC (2006).

17 Shang, T. et al. Hypoxia promotes differentiation of adipose-derived stem cells into endothelial cells through demethylation of ephrinb2. Stem Cell Res Ther 10, 133, doi:10.1186/s13287-019-1233-x (2019).

18 Bekhite, M. M. et al. Hypoxia, leptin, and vascular endothelial growth factor stimulate vascular endothelial cell differentiation of human adipose tissue-derived stem cells. Stem Cells Dev 23, 333–351, doi:10.1089/scd.2013.0268 (2014).

19 Podkalicka, P. et al. Hypoxia as a driving force of pluripotent stem cell reprogramming and differentiation to endothelial cells. Biomolecules 10, doi:10.3390/biom10121614 (2020).

20 Yuan, C. et al. Coculture of stem cells from apical papilla and human umbilical vein endothelial cell under hypoxia increases the formation of three-dimensional vessel-like structures in vitro. Tissue Eng Part A 21, 1163–1172, doi:10.1089/ten.TEA.2014.0058 (2015).

21 Salomon, C. et al. Exosomal signaling during hypoxia mediates microvascular endothelial cell migration and vasculogenesis. PLoS One 8, e68451, doi:10.1371/journal.pone.0068451 (2013).

22 Prado-Lopez, S. et al. Hypoxia promotes efficient differentiation of human embryonic stem cells to functional endothelium. Stem Cells 28, 407–418, doi:10.1002/stem.295 (2010).

23 Han, Y., Kuang, S. Z., Gomer, A. & Ramirez-Bergeron, D. L. Hypoxia influences the vascular expansion and differentiation of embryonic stem cell cultures through the temporal expression of vascular endothelial growth factor receptors in an arnt-dependent manner. Stem Cells 28, 799–809, doi:10.1002/stem.316 (2010).

24 Loughna, S., Yuan, H. T. & Woolf, A. S. Effects of oxygen on vascular patterning in tie1/lacz metanephric kidneys in vitro. Biochem Biophys Res Commun 247, 361–366, doi:10.1006/bbrc.1998.8768 (1998).

25 Tufro-McReddie, A. et al. Oxygen regulates vascular endothelial growth factor-mediated vasculogenesis and tubulogenesis. Dev Biol 183, 139–149, doi:10.1006/dbio.1997.8513 (1997).

26 Fajersztajn, L. & Veras, M. M. Hypoxia: From placental development to fetal programming. Birth Defects Res 109, 1377–1385, doi:10.1002/bdr2.1142 (2017).

27 Hemker, S. L., Sims-Lucas, S. & Ho, J. Role of hypoxia during nephrogenesis. Pediatric nephrology (Berlin, Germany) 31, 1571–1577, doi:10.1007/s00467-016-3333-5 (2016).

28 Keeley, T. P. & Mann, G. E. Defining physiological normoxia for improved translation of cell physiology to animal models and humans. Physiol Rev 99, 161–234, doi:10.1152/physrev.00041.2017 (2019).

29 Tufro, A. Vegf spatially directs angiogenesis during metanephric development in vitro. Dev Biol 227, 558–566, doi:10.1006/dbio.2000.9845 (2000).

30 Bernhardt, W. M. et al. Expression of hypoxia-inducible transcription factors in developing human and rat kidneys. Kidney Int 69, 114–122, doi:10.1038/sj.ki.5000062 (2006).

31 Takasato, M. et al. Kidney organoids from human ips cells contain multiple lineages and model human nephrogenesis. Nature 526, 564–568, doi:10.1038/nature15695 (2015).

32 van den Berg, C. W. et al. Renal subcapsular transplantation of psc-derived kidney organoids induces neo-vasculogenesis and significant glomerular and tubular maturation in vivo. Stem Cell Rep. 10, 751–765, doi:10.1016/j.stemcr.2018.01.041 (2018).

33 Geuens, T. et al. Thiol-ene cross-linked alginate hydrogel encapsulation modulates the extracellular matrix of kidney organoids by reducing abnormal type 1a1 collagen deposition. Biomaterials 275, 120976, doi:10.1016/j.biomaterials.2021.120976 (2021).

34 Schindelin, J. et al. Fiji: An open-source platform for biological-image analysis. Nat Methods 9, 676–682, doi:10.1038/nmeth.2019 (2012).

35 Schneider, C. A., Rasband, W. S. & Eliceiri, K. W. Nih image to imagej: 25 years of image analysis. Nat Methods 9, 671–675, doi:10.1038/nmeth.2089 (2012).

36 Xie, F., Xiao, P., Chen, D., Xu, L. & Zhang, B. Mirdeepfinder: A mirna analysis tool for deep sequencing of plant small rnas. Plant Mol Biol, doi:10.1007/s11103-012-9885-2 (2012).

37 Klingberg, A. et al. Fully automated evaluation of total glomerular number and capillary tuft size in nephritic kidneys using lightsheet microscopy. J Am Soc Nephrol 28, 452–459, doi:10.1681/ASN.2016020232 (2017).

38 Freeburg, P. B., Robert, B., St John, P. L. & Abrahamson, D. R. Podocyte expression of hypoxia-inducible factor (hif)-1 and hif-2 during glomerular development. J Am Soc Nephrol 14, 927–938, doi:10.1097/01.asn.0000059308.82322.4f (2003).

39 Vempati, P., Popel, A. S. & Mac Gabhann, F. Extracellular regulation of vegf: Isoforms, proteolysis, and vascular patterning. Cytokine & growth factor reviews 25, 1–19, doi:10.1016/j.cytogfr.2013.11.002 (2014).

40 Otrock, Z. K., Mahfouz, R. A., Makarem, J. A. & Shamseddine, A. I. Understanding the biology of angiogenesis: Review of the most important molecular mechanisms. Blood Cells Mol Dis 39, 212–220, doi:10.1016/j.bcmd.2007.04.001 (2007).

41 Michiels, C., Arnould, T. & Remacle, J. Endothelial cell responses to hypoxia: Initiation of a cascade of cellular interactions. Biochim Biophys Acta 1497, 1–10, doi:10.1016/s0167-4889(00)00041-0 (2000).

42 Haddad, J. J. & Harb, H. L. Cytokines and the regulation of hypoxia-inducible factor (hif)-1alpha. Int Immunopharmacol 5, 461–483, doi:10.1016/j.intimp.2004.11.009 (2005).

43 Abhinand, C. S., Raju, R., Soumya, S. J., Arya, P. S. & Sudhakaran, P. R. Vegf-a/vegfr2 signaling network in endothelial cells relevant to angiogenesis. J Cell Commun Signal 10, 347–354, doi:10.1007/s12079-016-0352-8 (2016).

44 Guan, F., Villegas, G., Teichman, J., Mundel, P. & Tufro, A. Autocrine vegf-a system in podocytes regulates podocin and its interaction with cd2ap. Am J Physiol Renal Physiol 291, F422–428, doi:10.1152/ajprenal.00448.2005 (2006).

45 Freeburg, P. B. & Abrahamson, D. R. Hypoxia-inducible factors and kidney vascular development. J Am Soc Nephrol 14, 2723–2730, doi:10.1097/01.asn.0000092794.37534.01 (2003).

46 Gerl, K. et al. Activation of hypoxia signaling in stromal progenitors impairs kidney development. Am J Pathol 187, 1496–1511, doi:10.1016/j.ajpath.2017.03.014 (2017).

47 Kobayashi, H. et al. Hypoxia-inducible factor prolyl-4-hydroxylation in foxd1 lineage cells is essential for normal kidney development. Kidney Int 92, 1370–1383, doi:10.1016/j.kint.2017.06.015 (2017).

48 Yousef Yengej, F. A., Jansen, J., Rookmaaker, M. B., Verhaar, M. C. & Clevers, H. Kidney organoids and tubuloids. Cells 9, 1326, doi:10.3390/cells9061326 (2020).

49 Shih, H. M., Wu, C. J. & Lin, S. L. Physiology and pathophysiology of renal erythropoietin-producing cells. J Formos Med Assoc 117, 955–963, doi:10.1016/j.jfma.2018.03.017 (2018).

50 Kim, B. S., Chen, J., Weinstein, T., Noiri, E. & Goligorsky, M. S. Vegf expression in hypoxia and hyperglycemia: Reciprocal effect on branching angiogenesis in epithelial-endothelial co-cultures. J Am Soc Nephrol 13, 2027–2036, doi:10.1097/01.asn.0000024436.00520.d8 (2002).

51 Bevan, H. S. et al. The alternatively spliced anti-angiogenic family of vegf isoforms vegfxxxb in human kidney development. Nephron Physiol 110, p57–67, doi:10.1159/000177614 (2008).

52 Eremina, V. & Quaggin, S. E. The role of vegf-a in glomerular development and function. Curr Opin Nephrol Hypertens 13, 9–15, doi:10.1097/00041552-200401000-00002 (2004).

53 Farina, A. R. et al. Hypoxia-induced alternative splicing: The 11th hallmark of cancer. J Exp Clin Cancer Res 39, 110, doi:10.1186/s13046-020-01616-9 (2020).

54 Dahan, S. et al. Vegfa’s distal enhancer regulates its alternative splicing in cml. NAR Cancer 3, zcab029, doi:10.1093/narcan/zcab029 (2021).

55 Roodink, I. et al. Development of the tumor vascular bed in response to hypoxia-induced vegf-a differs from that in tumors with constitutive vegf-a expression. Int J Cancer 119, 2054–2062, doi:10.1002/ijc.22072 (2006).

56 Cebe Suarez, S. et al. A vegf-a splice variant defective for heparan sulfate and neuropilin-1 binding shows attenuated signaling through vegfr-2. Cell Mol Life Sci 63, 2067–2077, doi:10.1007/s00018-006-6254-9 (2006).

57 Salton, M., Voss, T. C. & Misteli, T. Identification by high-throughput imaging of the histone methyltransferase ehmt2 as an epigenetic regulator of vegfa alternative splicing. Nucleic Acids Res 42, 13662–13673, doi:10.1093/nar/gku1226 (2014).

58 Peach, C. J. et al. Molecular pharmacology of vegf-a isoforms: Binding and signalling at vegfr2. International Journal of Molecular Sciences 19, 1264 (2018).

59 Stolz, D. B. & Sims-Lucas, S. Unwrapping the origins and roles of the renal endothelium. Pediatric nephrology (Berlin, Germany) 30, 865–872, doi:10.1007/s00467-014-2798-3 (2015).

60 Tahergorabi, Z. & Khazaei, M. A review on angiogenesis and its assays. Iran J Basic Med Sci 15, 1110–1126 (2012).

