## Supplementary figures for "Enhanced microvasculature formation and patterning in iPSC–derived kidney organoids cultured in physiological hypoxia"

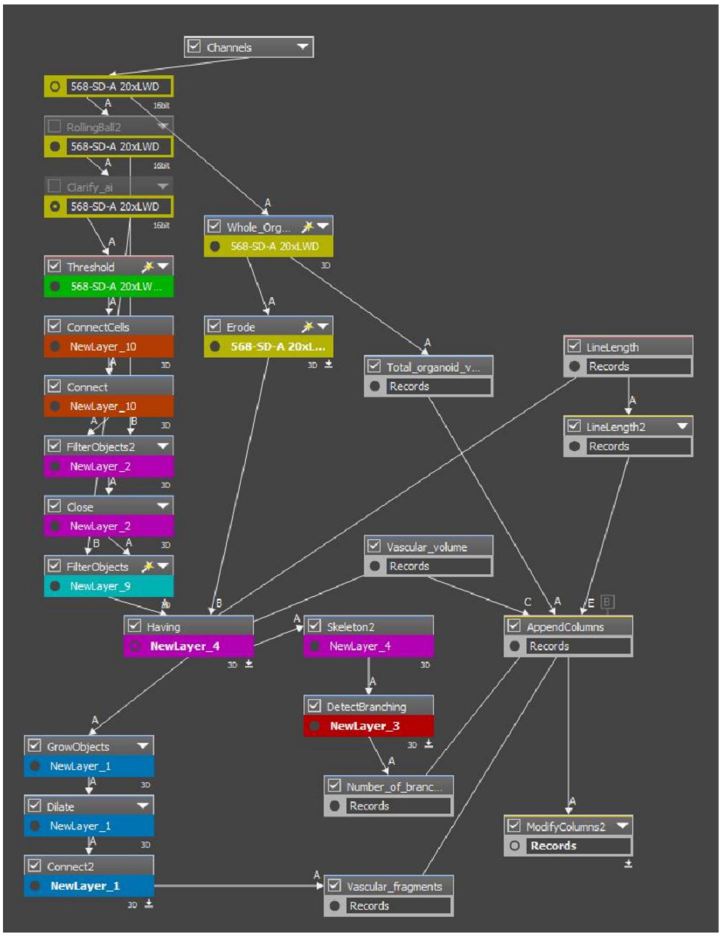


3D organoid view (raw)

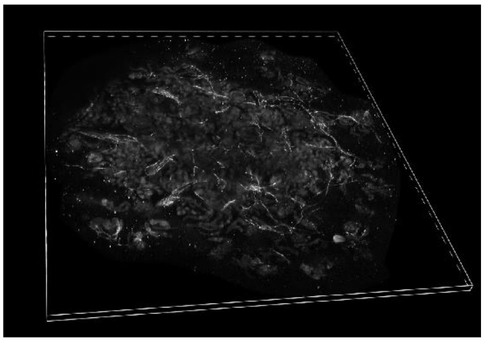

3D organoid segmentation

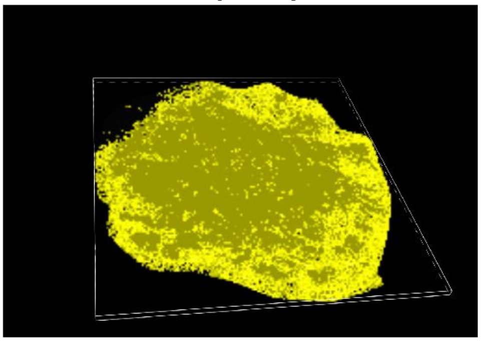

3D clarified CD31+ endothelial cells

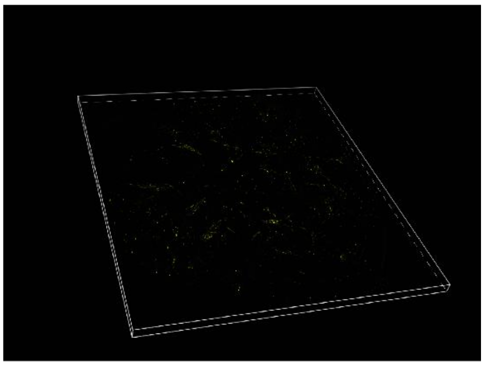

3D segmented CD31+ endothelial cells

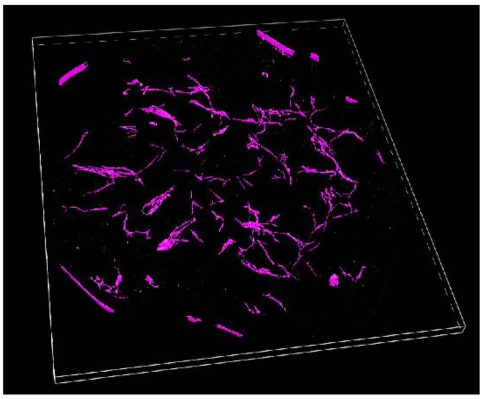

*Supplementary figure 1: Segmentation and quantification pipeline of the endothelial network in 3D. Detailed description can be found in the methods section. Briefly, after clarifying the image, the CD31 signal was thresholded, and after conservative filtering, the vessels were skeletonized and segmented.*

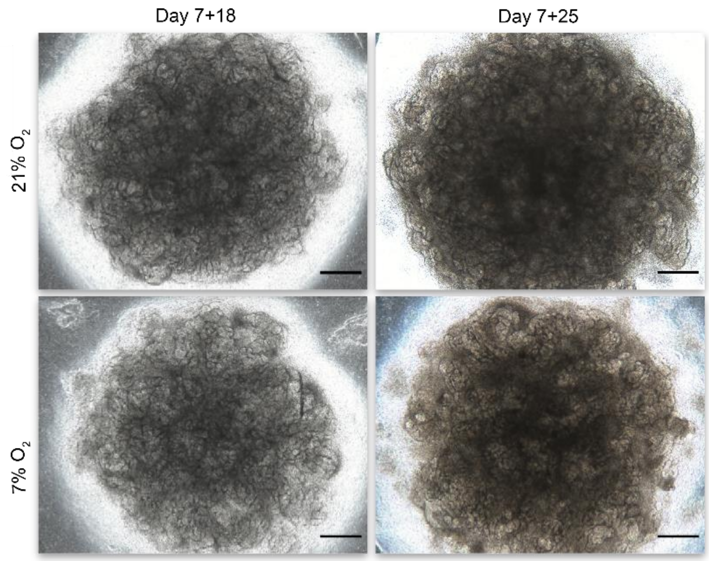


*Supplementary figure 2: Brightfield images of organoids in normoxia and hypoxia. Kidney organoids in 21% $O_2$ and 7% $O_2$ develop similar morphologies. Scale bars: 1 mm*

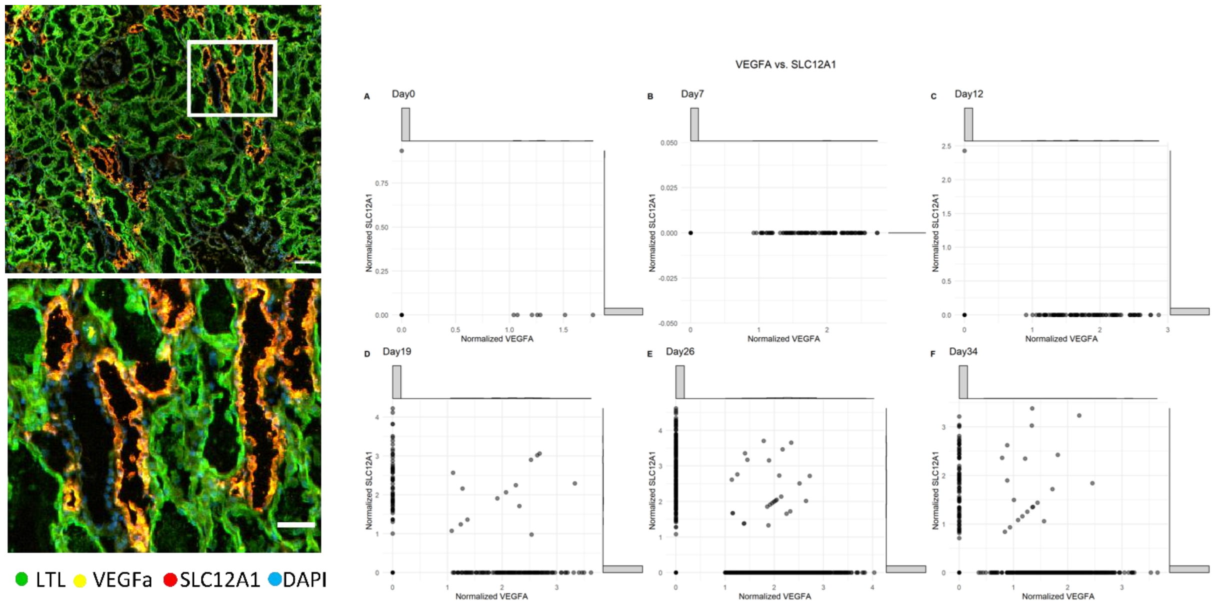


*Supplementary figure 3: Vascular endothelial growth factor alpha (VEGF-A) marker validation on adult human kidney and in single cell RNA sequencing datasets of kidney organoids. VEGF-A is co-expressed*

*with Solute carrier family 12 member 1 (SLC12A1) in both adult human kidney sections and single cell RNA sequencing data of Wu et al (2018). Scale bars: 50 µm*

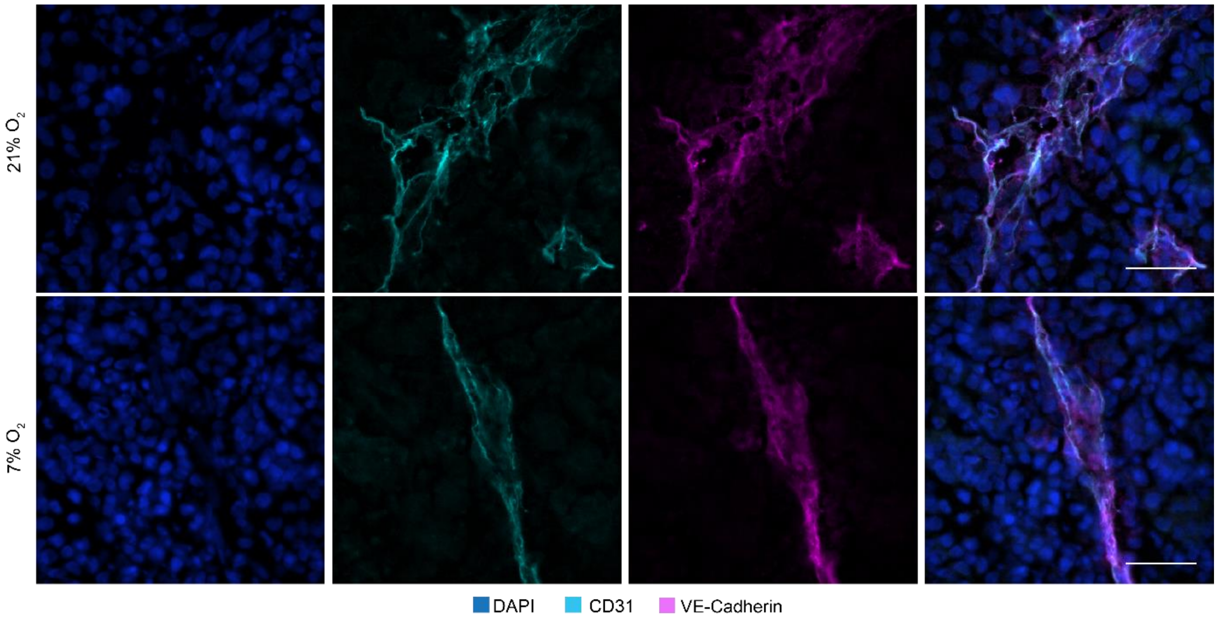


*Supplementary figure 4: CD31 and VE-cadherin expression at day 7+25 at 21% and 7% $O_2$. Co-expression of CD31 and VE-cadherin indicates maintenance of the endothelial phenotype at day 7+25 in culture in both 21% and 7% $O_2$. Scale bar: 30 µm*

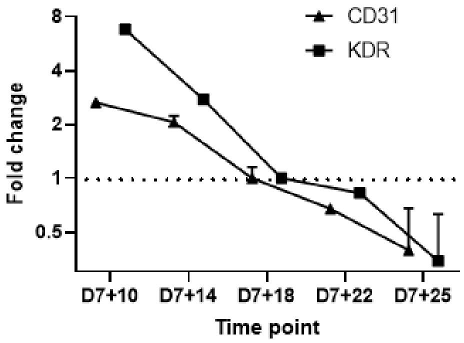


*Supplementary figure 5: Gene expression of CD31 and KDR is downregulated over time in 21% $O_2$ indicating a diminishing endothelial network.*

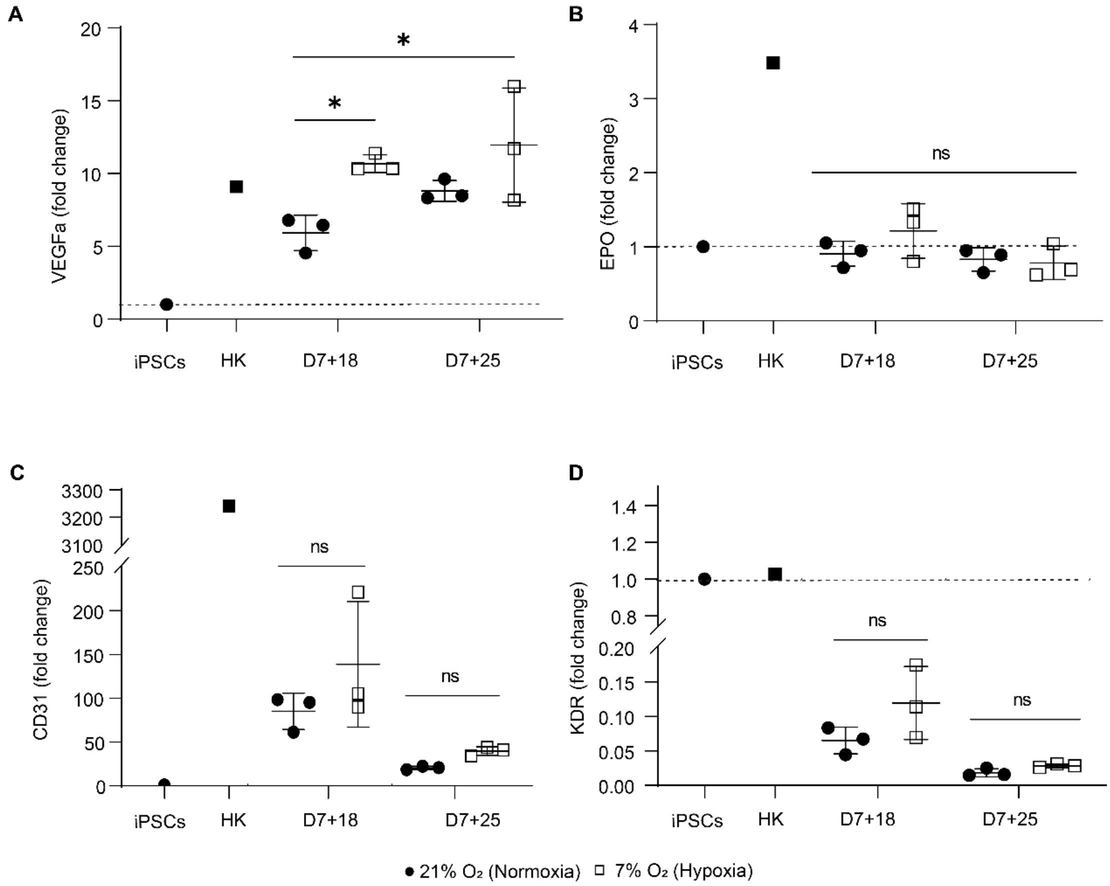


*Supplementary figure 6: Gene expression of VEGF-A (all variants), EPO, CD31 and KDR in 21% $O_2$ and 7% $O_2$ of a single experiment. VEGF-A mRNA is significantly upregulated at 7% $O_2$ (A). There is no significant difference between 21% $O_2$ and 7% $O_2$ in EPO (B), CD31 (C) and KDR (D) gene expression. Human kidney and iPSCs were plotted as reference. (n=3, N=1)*
